# Marseilleviruses biological properties and viral translation-associated proteins based on *in silico* tertiary structure

**DOI:** 10.1101/2024.06.22.600160

**Authors:** XIA Yucheng, Long Wang, Yutao Xu, Yue-Him Wong, Zhichao Zhou, ZHANG Rui

## Abstract

Marseilleviruses are a group of double-strand DNA viruses that infect amoeba within the *Nucleocytoviricota* phylum and are ubiquitous in water and soil globally. Here, we report four novel strains isolated from mangroves in Guangdong province, China, namely, futianmevirus, futianmfvirus, dashavirus str. E, and xiwanvirus. Viral particles presented about 220∼240 nm icosahedrally shaped capsids and were wrapped by membranes to form giant vesicles. Based on stability assays, viral particles were halotolerant and acid-tolerant, but sensitive to chloroform and high temperature, while giant vesicles conferred thermal and acid/alkaline resistance to particles. Genomics and phylogenetic analyses showed that the four strains formed divergent branches within different lineages of marseillevirus. Notably, to our knowledge, futianmevirus was the first reported marseillevirus lacking translation elongation factor EF-1alpha (EF1A). Our *in silico* analysis of marseillevirus coded translation-associated homolgs suggested their conserved functions. Additionally, we predicted at least four novel proteins that were structurally similar to components of the protozoan ribosome. Overall, not only our data comprehensively described the diversity of marseillevirus biological properties, but also proposed a new perspective on the giant virus translation system.

**Importance:** The family *Marseilleviridae* was the second reported family of giant viruses and distributed globally. In this work, we reported the four novel marseilleviruses isolated from saltwater samples of mangrove. Difference of biological properties between giant vesicles and viral particles revealed the environment fitness of marseillevirus.

On the other hand, sensitivity to chloroform indicated the importance of lipid components for viral infection. Additionally, our comparative genomics, phylogenetic analysis, and protein structure comparison revealed the diverse translation-associated gene sets of marseilleviruses. The prediction of ribosome components expands the knowledge about the giant viral translation-associated proteins, and will be helpful in future to reveal how giant viruses hijack the amoeba translation system.

## Introduction

The first described giant virus, *Acanthamoeba polyphaga* mimivirus, including the discovery of viral translation-associated proteins, has blurred some well-established virological concepts, irreversibly [1, 2]. Using similar isolation strategy on *Acanthamoeba*, the second described giant virus, Marseillevirus marseillevirus strain T19, was isolated from a water sample collected in a cooling tower in Paris, France [3]. It is the founder of a new viral family, officially recognized and named *Marseilleviridae* by the International Committee of Taxonomy of Viruses [4]. Since 2009, it has been shown that marseilleviruses are ubiquitous globally in water, soil, sediment, biospecimens and so on [5]. Based on the molecular phylogenetic analysis of the major capsid protein, the isolated members of marseilleviridae are currently classified into five lineages (A, B, C, D and E) [5].

The morphogenesis of marseillevirus particles vary in different lineages. Marseillevirus str. T19, a member of lineage A, has icosahedral virions with a diameter of approximately 250 nm with fibrils on the surface, while Lausannevirus, a member of lineage B, has smaller icosahedral viral particles without fibrils [3, 6].

Marseillevirus str. T19 is able to form giant vesicles containing various numbers of particles to stimulate amoeba phagocytosis and initiate virus replication cycle in the cytoplasm [7]. The giant vesicles have a higher efficacy and higher heat resistance than the particle suggesting that giant vesicles confer a physical advantage to marseillevirus [7]. However, the study of biological properties of giant viral particles under different environments was still limited.

The genomes of marseillevirus are circular double strand DNA with a size ranging from 348 to 404 kilobases (kb), and with a GC content varying between 42.9% and 44.8% [5]. It was hypothesized that the genome repertoire of marseillevirus was chimeric since eukaryotic homologs, like histones, and bacterial/ bacteriophage homologs, like MORN (Membrane Occupation and Recognition Nexus)-repeat-domain containing proteins, were common in marseilleviruses but barely in other giant viruses [3]. What’s more, there were four translation-associated genes, translation initiation factor SUI1 (SUI1), eukaryotic translation initiation factor 5 (eIF5), translation elongation factor EF-1alpha (EF1A) and eukaryotic peptide chain release factor 1 (eRF1) encoded by marseillevirus str. T19, and they were conserved among the members of *Marseilleviridae* [2, 3]. However, up to now, less than 40% of viral proteins were annotated, and there was a high proportion of uncharacterized proteins with no function assigned even using a search of three-dimensional structural homologs [8].

Here we reported the biological properties and genomic characterization of four new members of marseilleviridae, namely futianmevirus, futianmfvirus, dashavirus str. E and xiwanvirus. They were isolated from saltwater samples collected in mangroves in Guangdong province, China. Based on the results of phylogenetic analysis, futianmevirus and futianmfvirus formed divergent branches within the lineage A and close to the marseillevirus ancestor, and dashavirus str. E and xiwanvirus were close to other reported isolates in the lineage B. Notably, to our knowledge, futianmevirus was the first reported marseillevirus lacking EF1A. Based on tertiary structure homologs, the HHpred and alphafold annotated more translation-associated proteins, four of which were similar to the component proteins of protozoan ribosome complexes. Overall, our results suggested diverse marseilleviruses with different biological properties and translation-associated gene sets.

## Results

### Morphology and replication cycle

The morphology of the icosahedral viral particles observed using transmission electron microscopy (TEM) showed that the diameters of futianmevirus, futianmfvirus, dashavirus str. E and xiwanvirus particles were 240 ± 12 nm, 245 ± 15 nm, 236 ± 13 nm, and 226 ± 14 nm, respectively, and distributed in the cytoplasm at 2 hour post infection (h p.i.) before the lysis of cell (**Fig. 1A**).

**Fig. 1.**
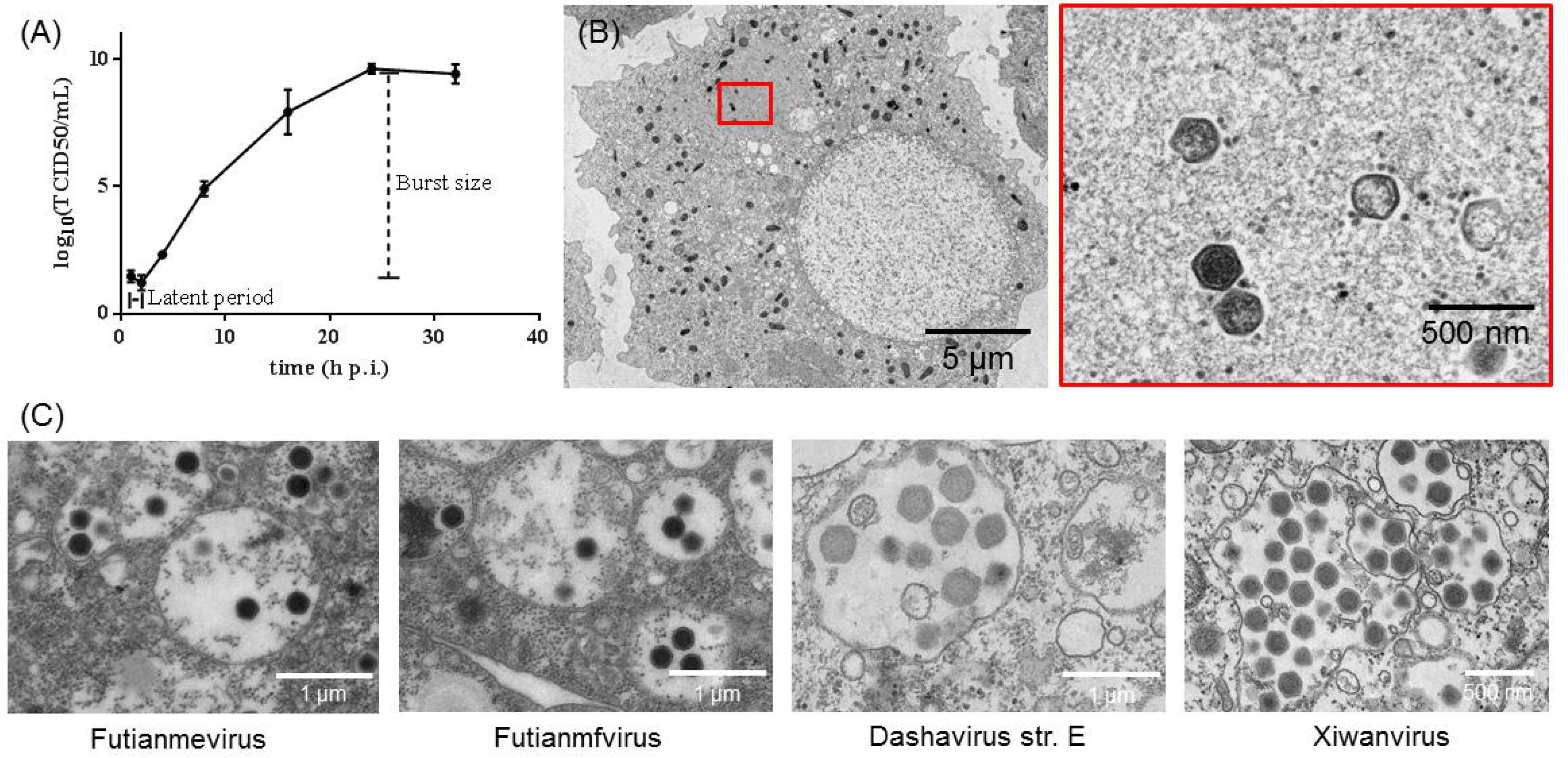
Virus replication and morphology of marseillevirus. (A) The viral replication logarithmic curve of futianmevirus. (B) Transmission electron micrographs of *A. castellanii* cells infected with futianmevirus showing the viral factory at 2 h p.i. (C) and different viral vesicles containing mature particles at 48 h p.i.

The one-step growth curve of futianmevirus, futianmfvirus, dashavirus str. E and xiwanvirus showed that latent period were approximately 4 hours (**Fig. 1B**, **Fig. 10**). The burst sizes is the total number of infectious particles released, which were treated with lysis buffer, at the end of one cycle of replication/number of infected amoebae.

The burst sizes of futianmevirus, futianmfvirus, dashavirus str. E and xiwanvirus, were 2,336 ± 407 particles/cell, 2,563 ± 822 particles/cell, 4,760 ± 650 particles/cell, and 4,759 ± 788 particles/cell, respectively.

### Viral particles released and chloroform sensitivity

Viral particles were wrapped by membranes, which were originated from the endoplasmic reticulum, as a giant vesicle to efficiently infect amoebae [7]. To release viral particles from giant vesicles, lysis buffer and chloroform were chosen to treat purified viruses separately. The lysis buffer had been described in the literature [7], while chloroform was served as an indication of structural lipid components of infection. The end-point dilution assays revealed that more infectious particles were released from dashavirus or xiwanvirus vesicles than that of futianmevirus or futianmfvirus indicating that the number of particles in vesicles varied in different marseilleviruses (**Fig. 1C**, **Fig. 2A**).

**Fig. 2.**
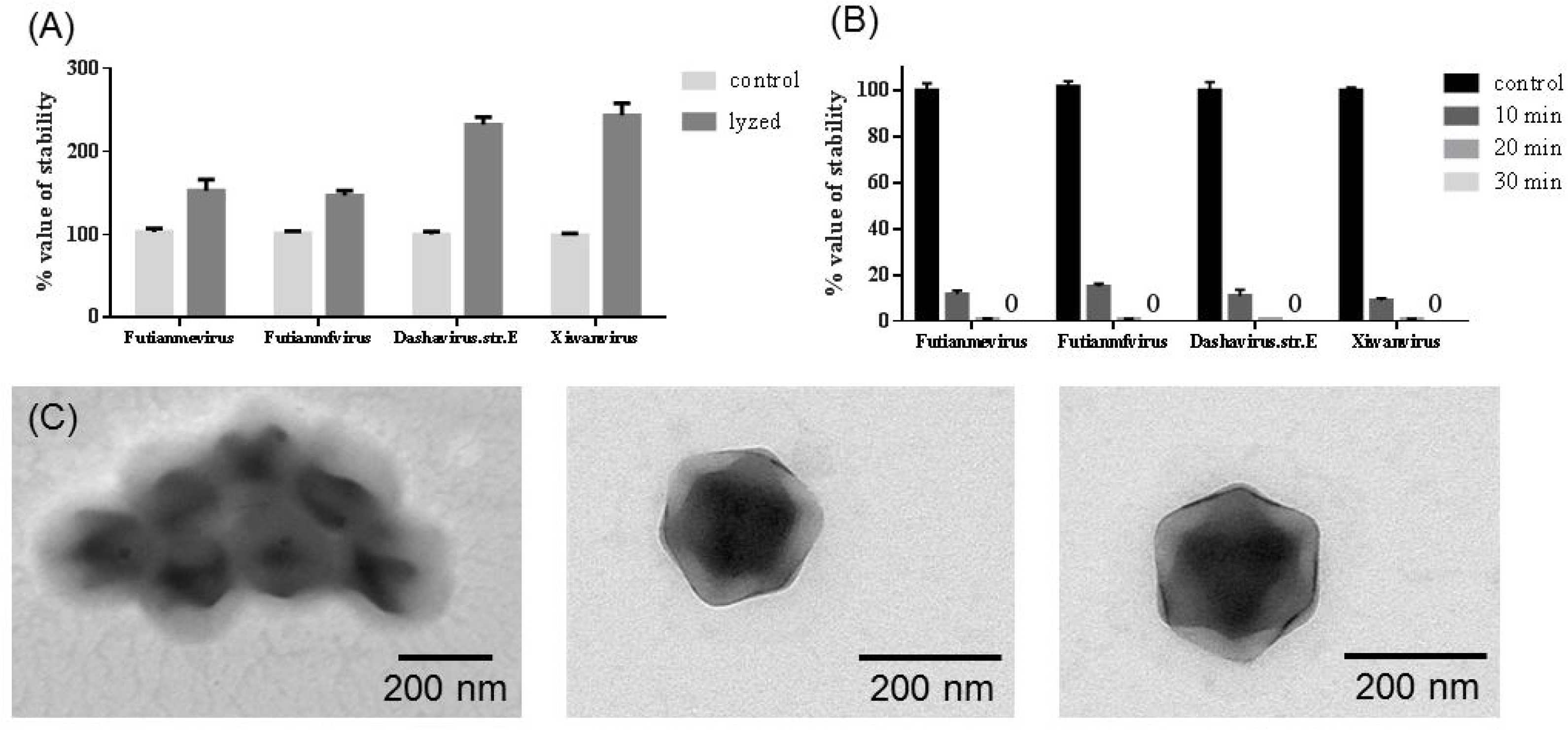
Effect of organic solvents on the infectivity of marseilleviruses. (A) Histogram of the percentage of infectious vesicles/particles after treatment with a lysis buffer. (B) Histogram of the percentage of infectious particles after treatment with chloroform. (C) Negative staining of a vesicle, a particle, and a particle chloroform-treated.

Interestingly, we observed that all marseilleviruses were highly sensitive to chloroform treatment indicating that lipid components were very important for marseillevirus infection (**Fig. 2B**). However, there was no morphological change between viral particles chloroform-treated and untreated (**Fig. 2C**), since chloroform sensitivity alone did not show the presence of lipid on the surface of viral particles [10].

### Stability of infectivity under different treatments

Considering the fact that mangrove habitats are characterized by high salinity, siltation, tidal surges, and flooding and waterlogged conditions [11], we tested the impacts of salinity, pH and temperature on the stability of giant vesicles and viral particles.

The incubation experiments of giant vesicles and viral particles in different concentrations of sodium chloride showed that all of them were halotolerant (**Fig. 3A**).

**Fig. 3.**
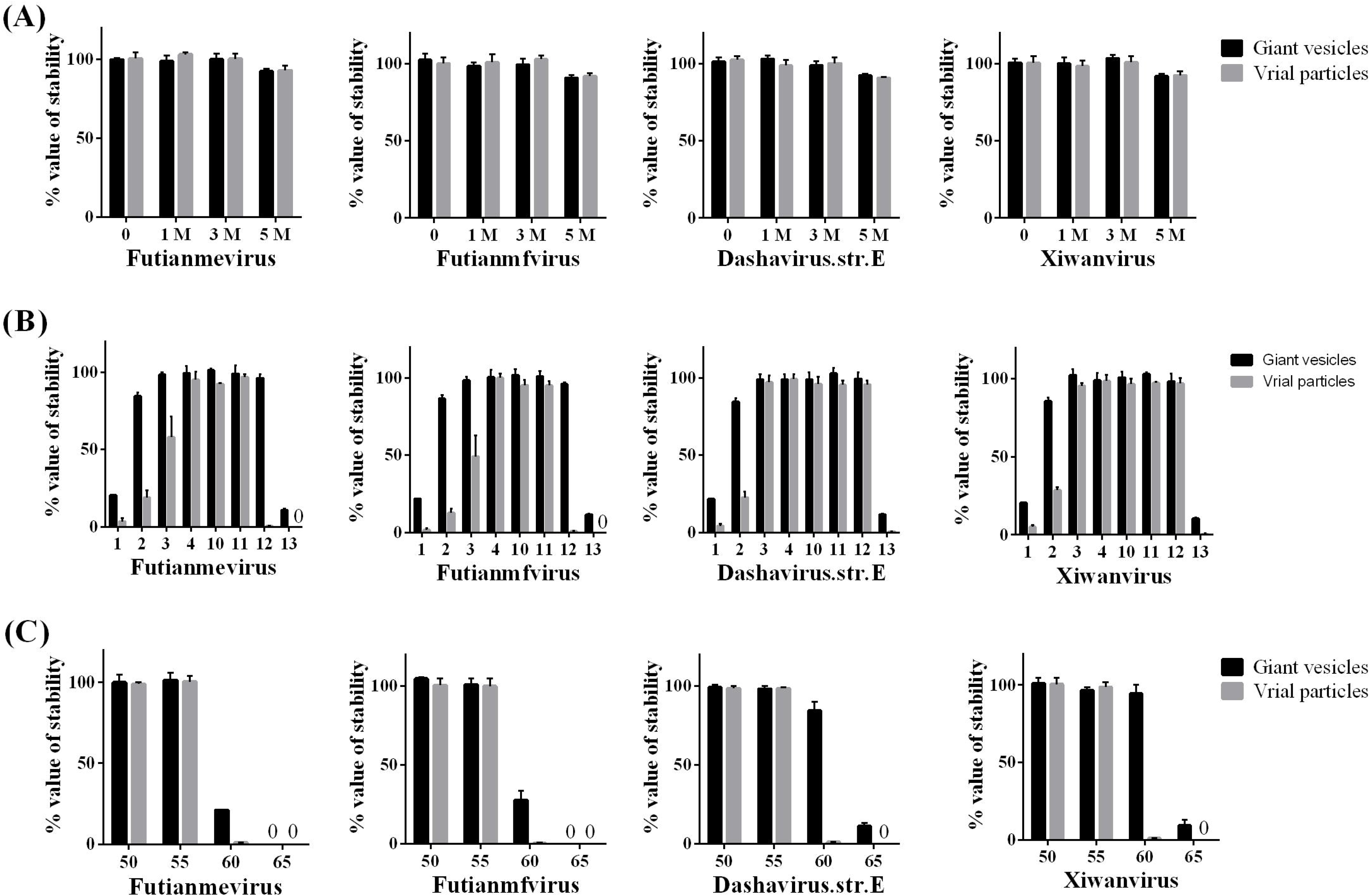
Biological properties of marseilleviruses. (A) Histogram of the percentage of infectious vesicles/particles after treatment with salt. (B) Histogram of the percentage of infectious vesicles/particles after treatment at different pH values. (C) Histogram of the percentage of infectious vesicles/particles after treatment at different temperatures. Marseillevirus titers were determined using the end-point dilution method. Data points represent the mean values standard deviations (SD) from three replicate experiments.

We dialyzed giant vesicles and viral particles against different sodium phosphate buffer solutions ranging in pH from 1 to 13. The incubation experiments showed that giant vesicles displayed a broad stability range with its infectivity maintained above 80% at pH 2 to 12, whereas, viral particles had a relatively narrow pH optimum of pH 3 to 11 of futianmevirus/ futianmfvirus, and pH 3 to 12 of dashavirus str. E/ xiwanvirus (**Fig. 3B**).

The thermal stability tests showed that viral particles were stable at 55°C for 1 h while giant vesicles were stable at 60°C for 1 h (**Fig. 3C**). Our results were consistent with the previous report that the viral vesicles were significantly more resistant to the thermal stress than the particles [7].

### Genomic content and phylogenetic analysis

Illumina sequencing and sequence assembly produced a single contig for each new marseillevirus genome, with size ranging from 360,239 to 372,628 bp (**Table 1**), which was in line with the genome sizes of reported marseilleviruses. The GC contents of the four marseilleviruses ranged from 42.26% to 45.25%, which were concordant with reported marseilleviruses. The numbers of predicted protein genes varied between 511 for futianmevirus and 523 for futianmfvirus (**Table 1**). A total of 2,064 proteins ranging from 29 to 2,065 amino acids (AA), with an average length of 221, were predicted. We found no evidence for the presence of tRNA and aminoacyl tRNA synthetase genes. The average distance between consecutive genes varied between 48-nt for dashavirus str. E and 56-nt for futianmevirus, resulting in an average coding density of 93.20%.

**TABLE 1.** Statistics of genome assemblies.

| Virus | Futianmevirus | Futianmfvirus | Dashavirus str. E | Xiwanvirus |
| --- | --- | --- | --- | --- |
| No. reads | 347912*2 | 781055*2 | 453751*2 | 1204782*2 |
| Ave. size / bp | 148 | 147 | 150 | 150 |
| No. contigs | 1 | 1 | 1 | 1 |
| Bases in all contigs / bp | 372,628 | 371,047 | 360,239 | 369,462 |
| N rate* / % | 0 | 0 | 0 | 100 |
| Coverage / % | 754 | 1,225 | 1,850 | 781 |
| GC content / % | 45.25 | 42.26 | 42.89 | 42.78 |
| No. CDS predicted | 511 | 523 | 511 | 519 |
| Intergene distance / nt | -60 ~ 580 | -54 ~ 516 | -111 ~ 468 | -111 ~ 446 |
| Coding density / % | 93.01 | 92.77 | 93.79 | 93.24 |
| GeneBank accession | OR157982.1 | OR343188.1 | OR343189.1 | MT663336.1 |

\*N rate: percentage of uncertain nucleotide.

To estimate the phylogenetic position of four marseilleviruses in *Marseilleviridae*, molecular phylogenetic analysis was performed based on NCLDV core genes, including major capsid protein, DNA polymerase delta catalytic subunit, D5-like primase-helicase, transcription factor S-II, and virion packaging ATPase [12]. The concatenated tree of five genes indicated that futianmevirus and futianmfvirus were grouped into lineage A, whereas dashavirus str. E and xiwanvirus were grouped into lineage B (**Fig. 4**).

**Fig. 4.**
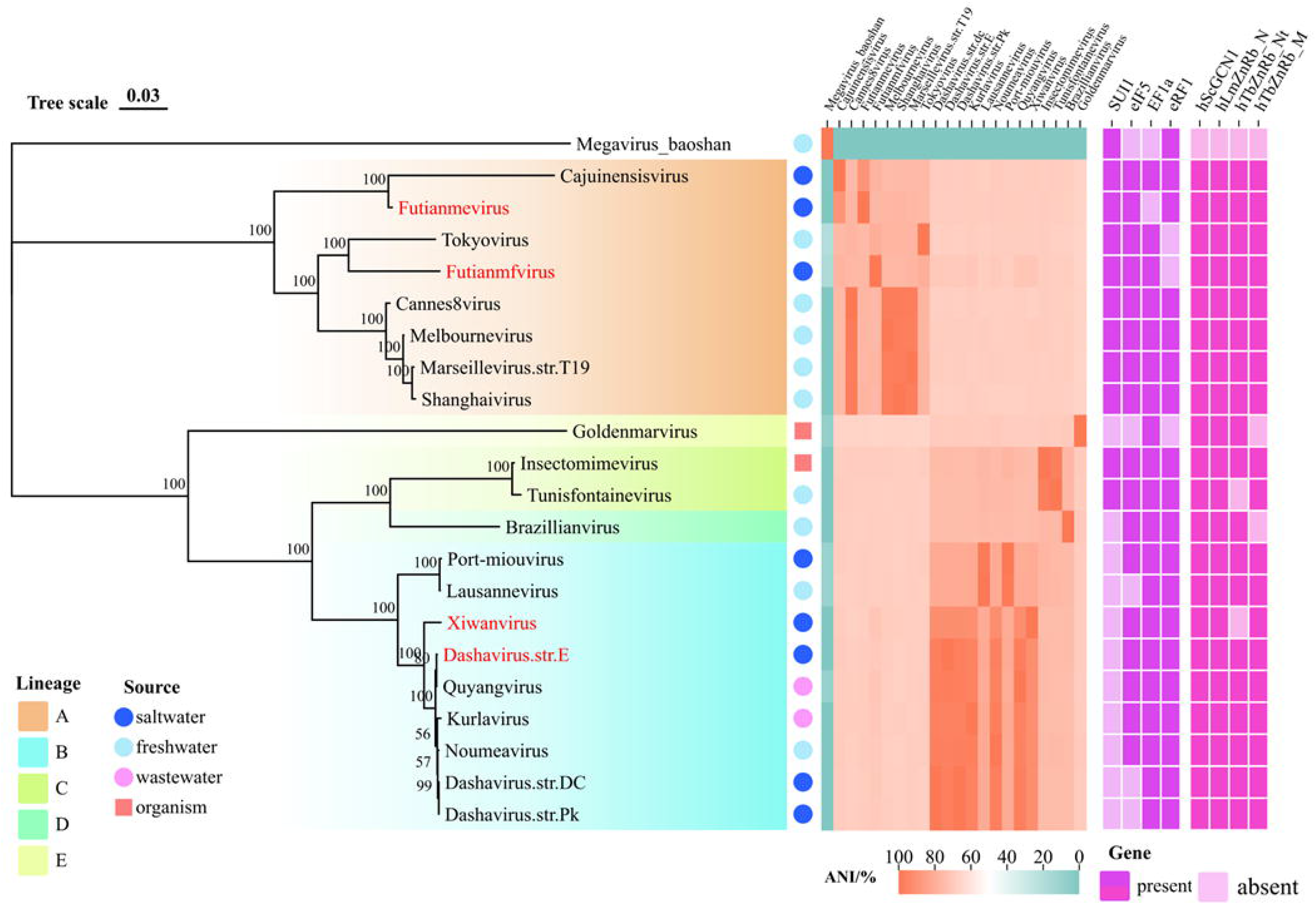
Molecular phylogenetic analysis based on the amino acid sequences of the concatenated NCLDV core genes. Numbers on each branch represented percentage of trees containing that branch. Background colors represented different lineages of viruses. Solid symbols adjacent to virus names represented environmental sources. The middle panel showed average nucleotide identity of whole viral genomes mapped to each other. The right panel showed the presence or absence of viral translation-associated genes, in which previous reported genes were in violet and putative ones found in this study were in pink.

### Comparative genomics and pan-/core-genome analysis

The average nucleotide identity (ANI) analysis of whole genome were performed to compare the nucleotide sequence divergence of marseilleviruses. The ANI of futianmevirus compared with cajuinensisvirus was 88.31% with the alignment percentage of 83.83 %, and that of futianmfvirus with tokyovirus was 79.50% (71.06%). Meanwhile, The ANIs of dashavirus str. E and xiwanvirus compared with kurlavirus were 96.10% (87.63%) and 90.87% (85.81%), respectively (**Fig. 4**).

Combining the previous results of phylogenetic analysis, we found that futianmevirus and futianmfvirus formed divergent branches within the lineage A and close to the viral ancestor, dashavirus str. E and xiwanvirus were close to other reported isolates in the lineage B.

A total of 10,987 predicted proteins, which were extracted from 21 isolated marseilleviruses, constituted a pangenome encompassing 900 clusters of orthologous proteins and 821 strain-specific proteins (**Fig. 5A**). Thirty eight clusters of orthologous proteins were involved in the pangenome with addition of four novel marseillevirus. The numbers of strain-specific proteins of futianmevirus, futianmfvirus, dashavirus str. E and xiwanvirus were 33, 43, 22 and 27, respectively. The core genome, containing at least one protein of each virus, included 169 clusters corresponding to 3,580 predicted proteins and represented 33% of the pangenome (**Fig. 5B**).

**Fig. 5.**
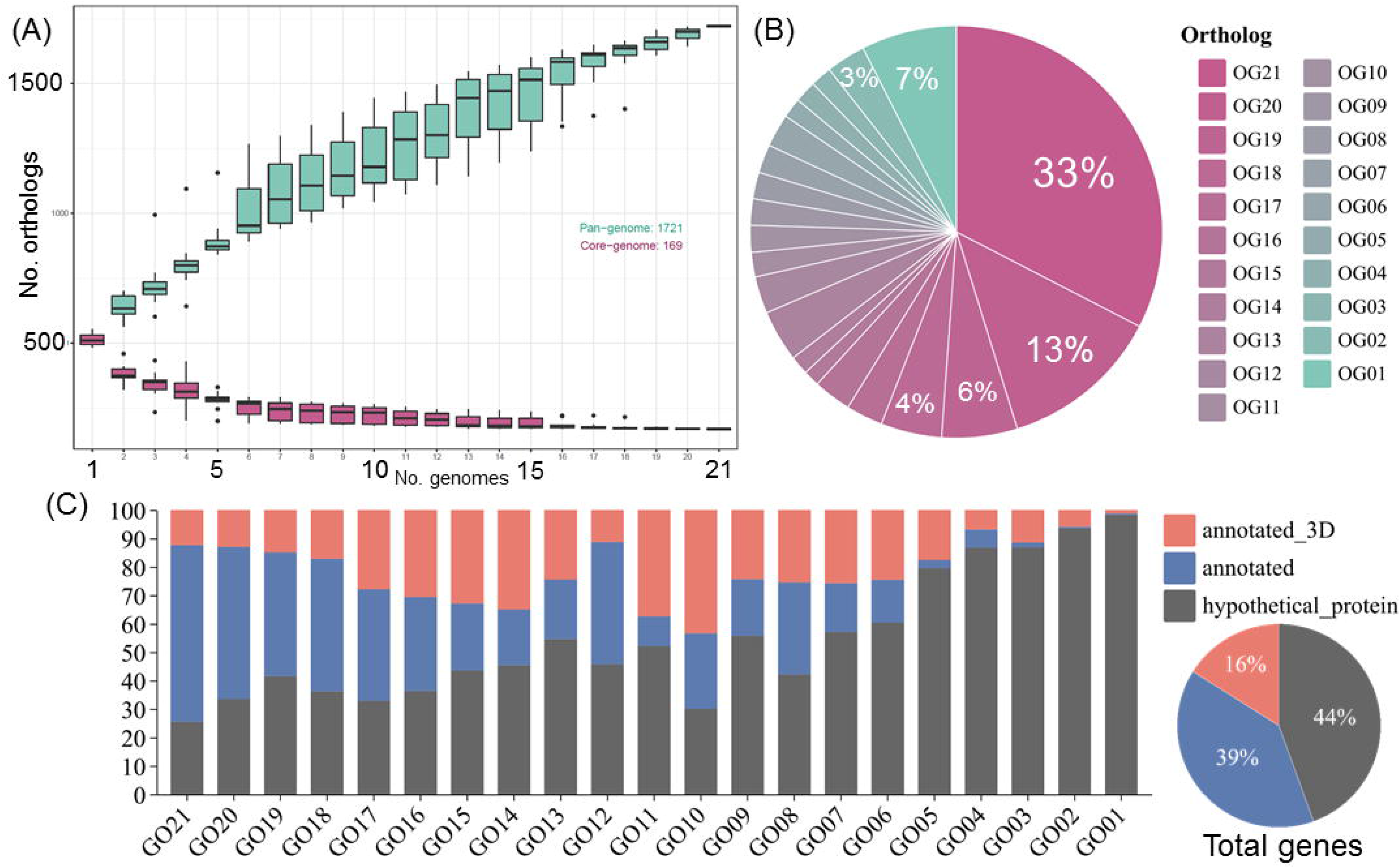
Orthology analysis of marseilleviruses. (A) The accumulation box curves represented pan-genomes and core-genomes of marseilleviruses. (B) The pie plot represented the percentage of viral orthologous genes in different gene groups, in which OG21 indicated the orthologous genes found in 21 strains of marseillevirs, OG20 for 20 strains and so forth. (C) The accumulation histogram and the pie plot represented the percentage of annotated genes in GenBank, ones in this study using structure-based prediction softwares with high confidence, and hypothetical proteins, the function of which could not be assigned using existing software.

Considering a high proportion of viral genes corresponded to hypothetical proteins without significantly similar sequences and function assigned, we applied protein structure prediction software, HHpred and alphafold2 for annotation of hypothetical proteins. Since functions were already known for a substantial proportion of proteins resident in pdb70 database, we uncovered structural homologs for ∼16% of viral proteins (**Fig. 5C**). Impressively, more translation-associated proteins were identified for viral hypothetical proteins (**Fig. 9**).

### Identification of reported translation-associated proteins

The results of amino acid residue alignments of EF1A and eRF1 were consistent with lineage classification and indicated their conservation (**Fig. 11, 12**). Our inspection of the nucleotide alignments showed that viral translation-associated genes were not located in highly conserved regions (**Fig. 6A**). Notably, EF1A, which was identified in all reported marseilleviruses, was absent in futianmevirus, and eRF1 was absent in futianmfvirus (**Fig. 4**). The absence of EF1A in futianmevirus or eRF1 in futianmfvirus was verified by PCR, respectively (**Fig. 6B**). The arrangements of orthologue proteins around EF1A or eRF1 reflected complex gene recombination events between different lineages and conservation within a lineage (**Fig. 6C**).

**Fig. 6.**
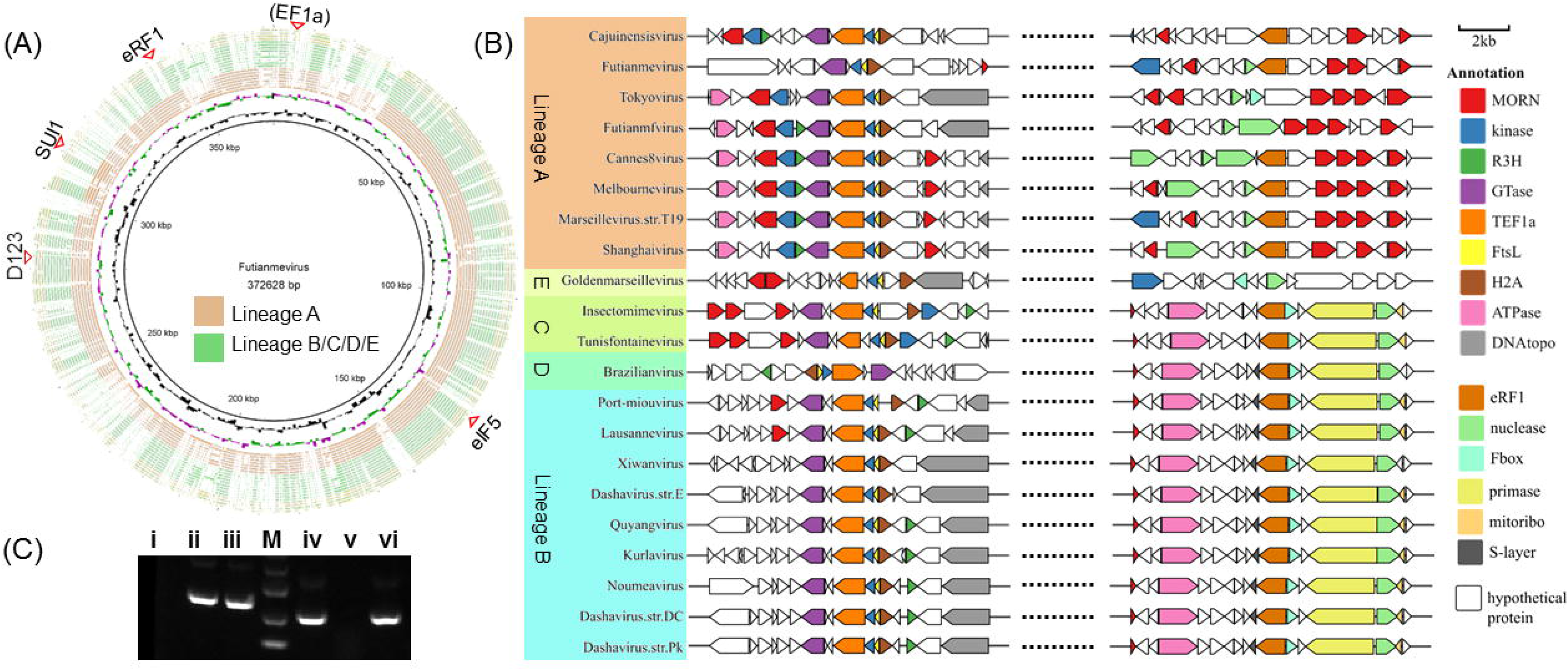
(A) Circular map of the futianmevirus chromosome. Rings starting from outer to innermost corresponded to (i) the complete genomic comparisons against 21 strains of marseillevirs with different colors representing lineages; (iii) GC skew for 1000-bp window and 500-bp step with skew+ in purple and skew-in green; (ii) GC content with higher or lower GC% compared with whole genome average. (B) Agarose gel electrophoresis analysis of PCR products. Lane i to iii: PCR products using a pair of primer targeting EF1A for futianmevirus, futianmfvirus and dashavirus str. E, respectively; Lane iv to vi: PCR products using a pair of primer targeting eRF1 for futianmevirus, futianmfvirus and dashavirus str. E, respectively. (C) Gene clusters of marseilleviruses displayed the relative size and direction of open reading frames near EF1A or eRF1. The scale is in kilobases. Abbreviations for labeled features are as follows: MORN, MORN-repeat-domain containing protein; R3H, R3H domain containing protein; FtsL, Cell division protein FtsL; H2A, histone H2A; DNAtopo, DNA topoisomerase II; Fbox, F-box domain containing protein; mitoribo, components of mitochondrial ribosome; S-layer, S-layer glycophorin.

**Fig. 7.**
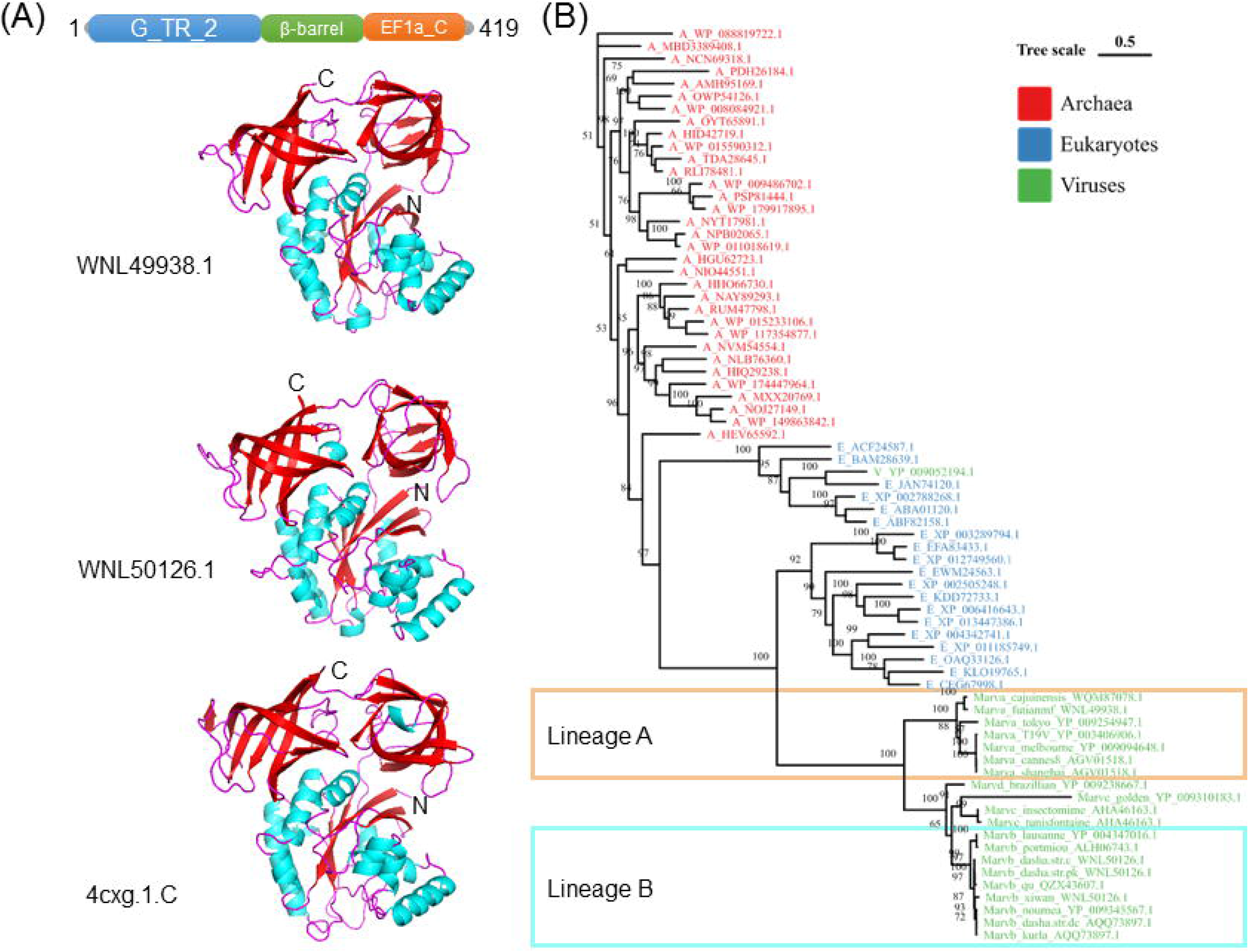
(A) Schematic representation of futianmfviral EF1A with the annotation of representative domains. Predicted structures of the futianmfviral EF1A, dashaviral EF1A, and mammalian EF1A. Abbreviations for labeled features are as follows: G_TR_2, translational type guanine nucleotide-binding domain; EF1A_C, beta-barrel domain that is found C-terminal of eEF1A. The predicted secondary structure were denoted in red for sheets, cyan for helixes, and pink for loops, respectively. GMQE score is expressed as a number between 0 and 1, reflecting the expected accuracy of a model built with the alignment and template, normalized by the coverage of the target sequence. (B) Molecular phylogenetic analysis based on the amino acid sequences of the EF1A genes. A color code was used to represent taxonomic groups, viruses in blue, archaea in red, and eukaryotes in green.

**Fig. 8.**
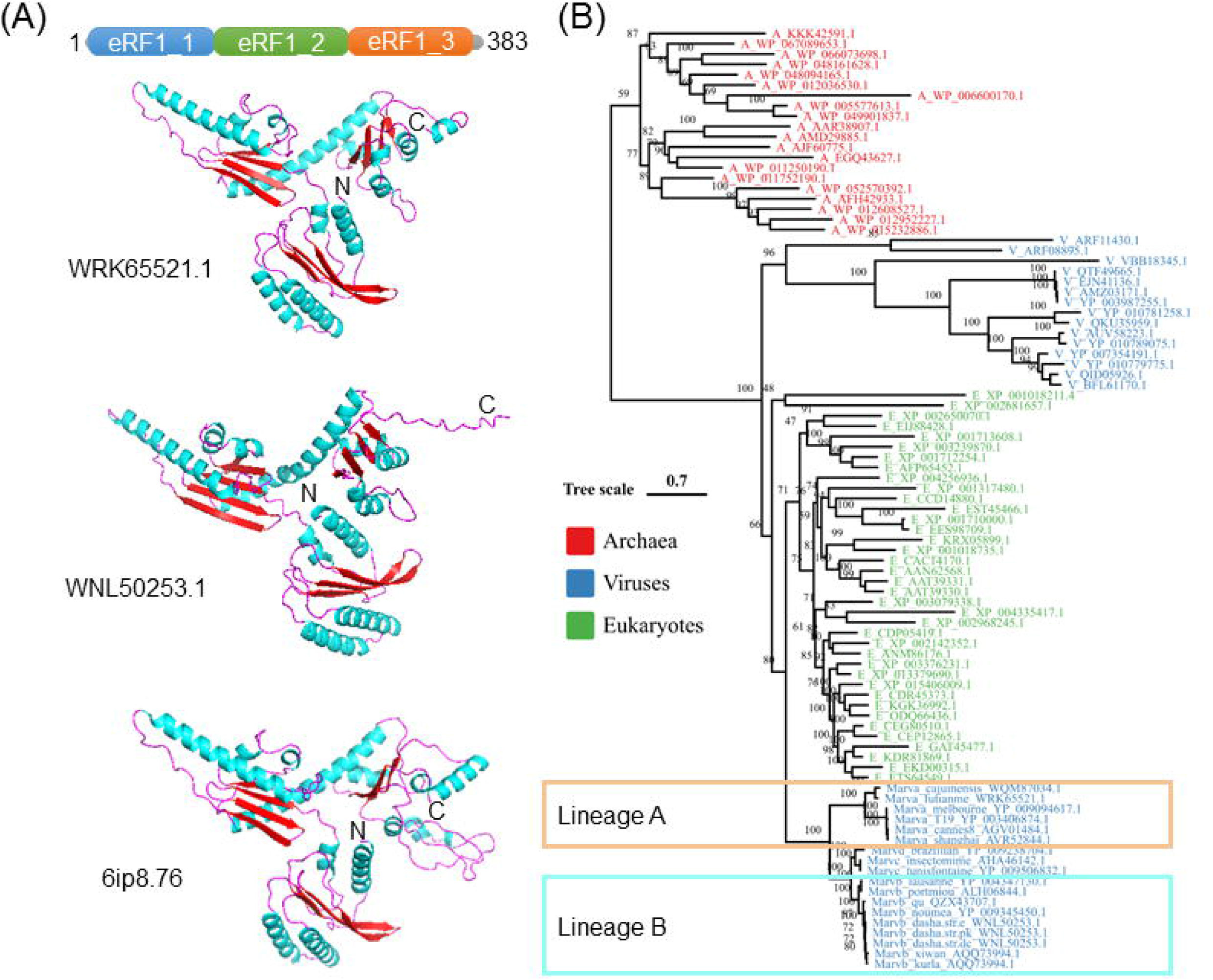
(A) Schematic representation of futianmeviral eRF1 with the annotation of representative domains. The overall shape and dimensions of eRF1 resembled a tRNA molecule with domains eRF1_1, eRF1_2, and eRF1_3 corresponding to the anticodon loop, aminoacyl acceptor stem, and T stem of a tRNA molecule, respectively. Predicted structures of the futianmeviral eRF1, dashaviral eRF1, and mammalian eRF1. The predicted secondary structure were denoted in red for sheets, cyan for helixes, and pink for loops, respectively. GMQE score is expressed as a number between 0 and 1, reflecting the expected accuracy of a model built with the alignment and template, normalized by the coverage of the target sequence. (B) Molecular phylogenetic analysis based on the amino acid sequences of the eRF1 genes. A color code was used to represent taxonomic groups, viruses in blue, archaea in red, and eukaryotes in green.

**Fig. 9.**
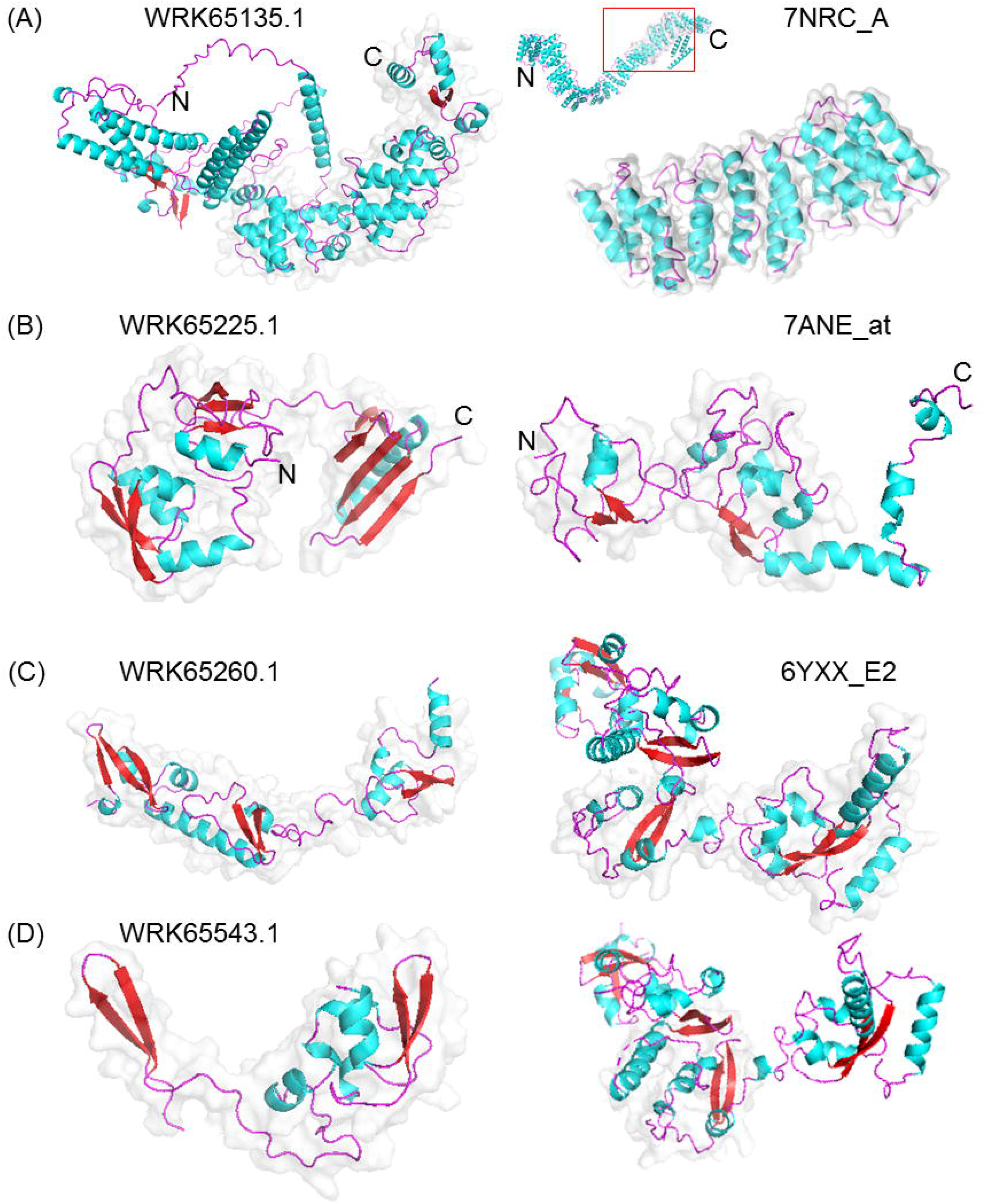
Predicted structures of new viral translation-associated proteins and templet proteins. (A) MarFTME_090, (B) MarFTME_180, (C) MarFTME_215 and (D) MarFTME_498. The predicted secondary structure were denoted in red for sheets, cyan for helixes, and pink for loops, respectively. The aligned tertiary structures were highlighted with grey outlines.

**Fig. 10.**
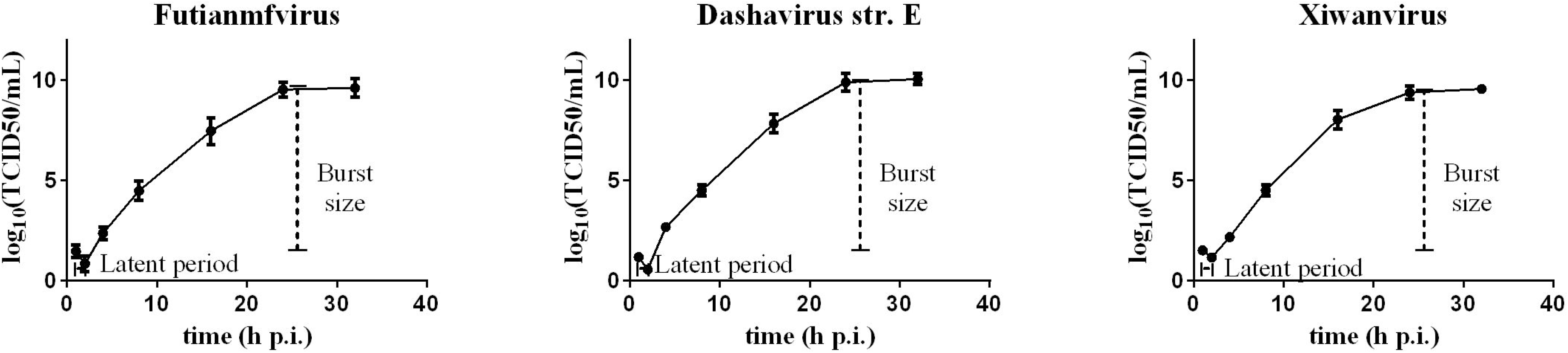
The viral replication logarithmic curves of futianmfvirus, dashavirus str. E, and xiwanvirus.

**Fig. 11.**
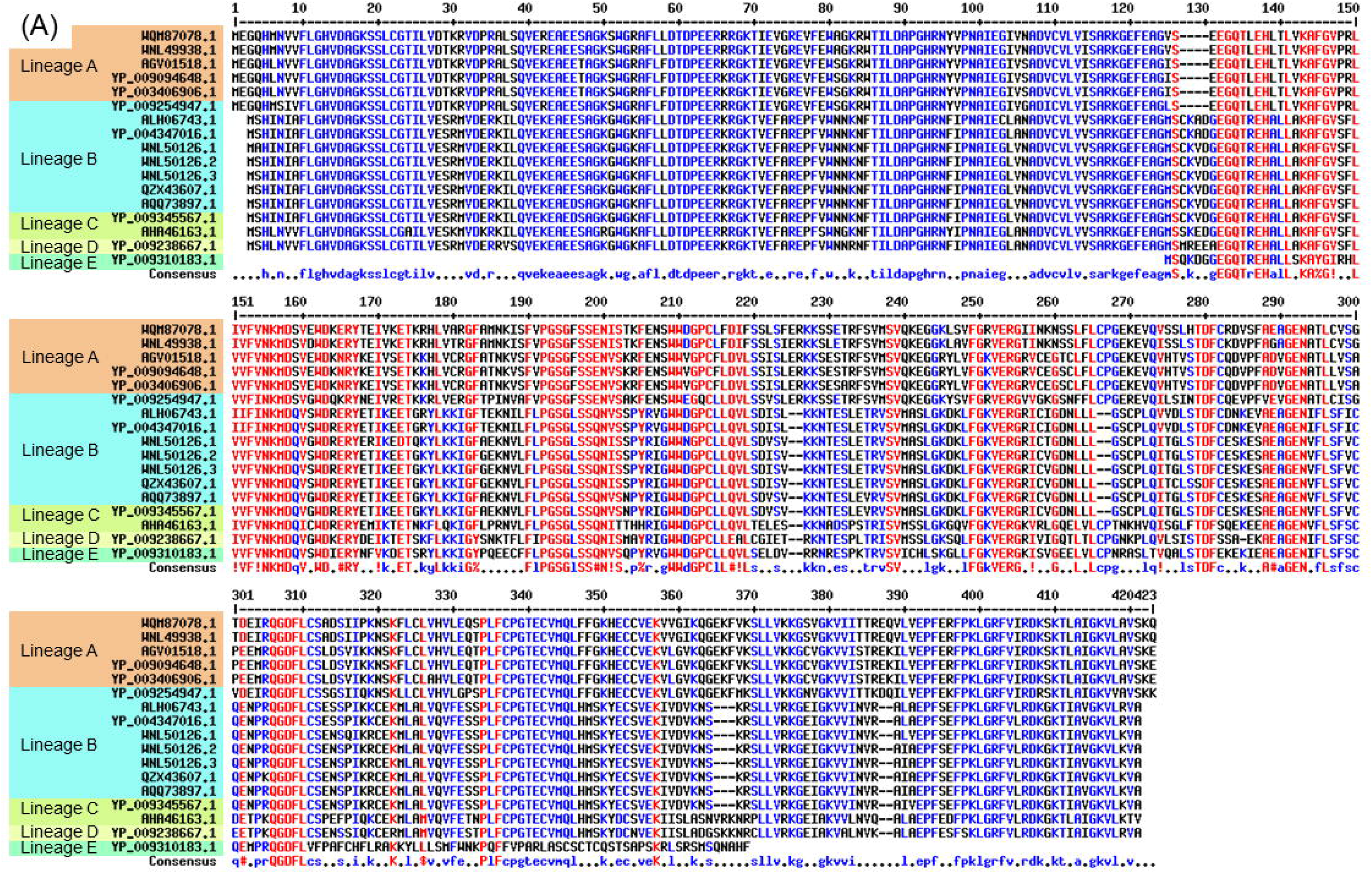
Multiple sequence alignments of viral EF1A using MUSCLE alignments. The high consensus (≥ 90%) residues was in red, low consensus (≤ 50%) residues was in blue, and neutral residues was in black.

**Fig. 12.**
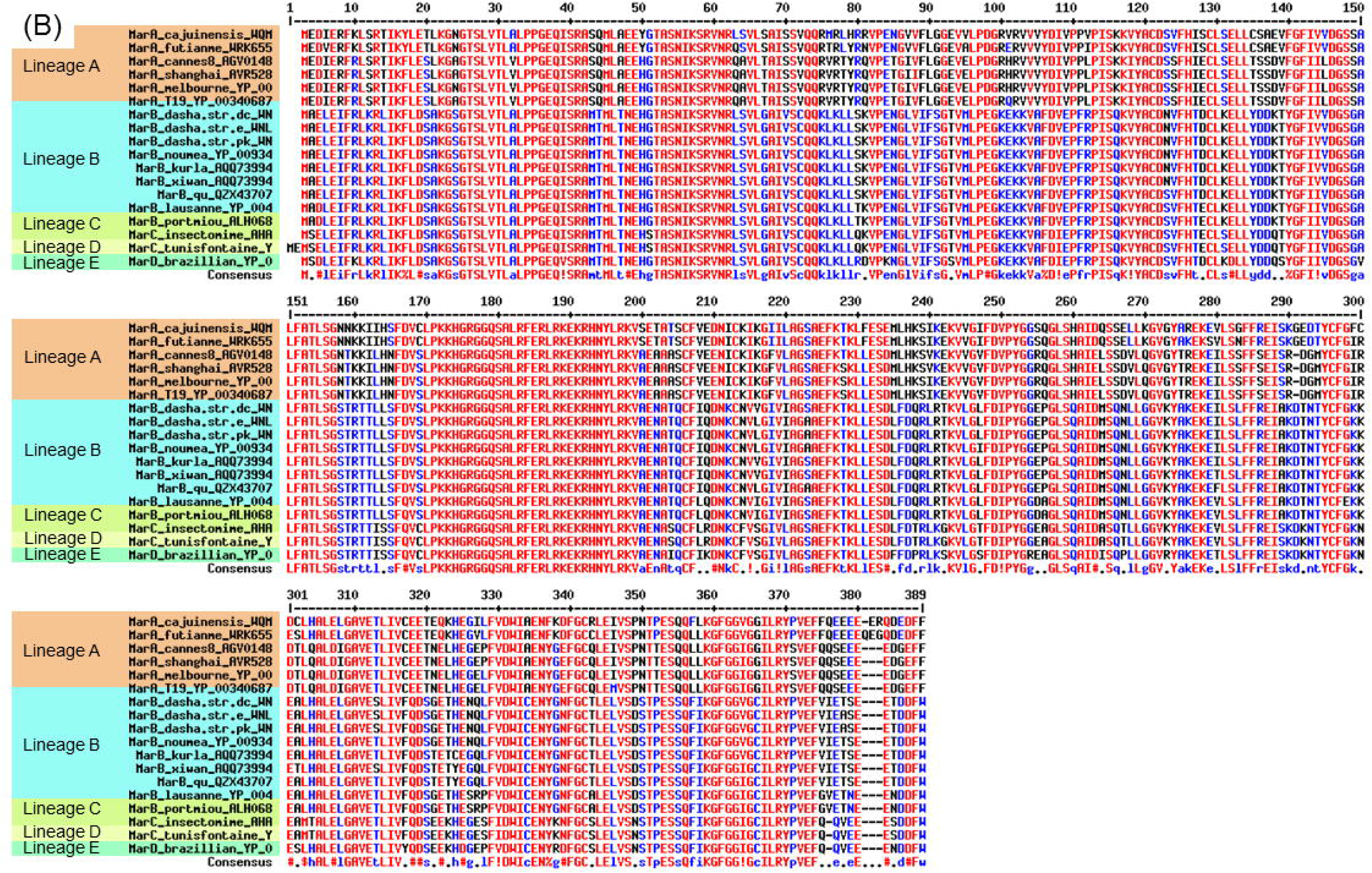
Multiple sequence alignments of viral eRF1 using MUSCLE alignments. The high consensus (≥ 90%) residues was in red, low consensus (≤ 50%) residues was in blue, and neutral residues was in black.

Besides, SUI1 was absent in marseillevirus of lineage B and goldenmarseillevirus (**Fig. 4**). The absence of reported translation-associated proteins in different marseilleviruses indicated the diverse translation-associated gene sets.

Based on conserved protein domain annotation and pair-wise structure comparison, viral EF1A and eRF1 were found to be similar to the eukaryotic ones. For instance, futianmfvirus EF1A (MarFTMF_422, GenBank: WNL49938.1) and dashavirus EF1A (MarDSR_087, WNL49938.1) had high scores with rabbit EF1A (4cxg.1.C) (**Fig. 7A**). Similarly, futianmevirus eRF1 (MarFTME_476, WRK65521.1) and dashavirus eRF1 (MarDSR_214, WNL50253.1) had high scores with anthropoid eRF1 (6ip8.76.A) (**Fig. 8A**). Therefore, the highly structural similarity indicating conserved function among eukaryotic and viral EF1A and eRF1. In addition, the results of phylogenetic analyses were compatible reported findings that giant viral translation factors affiliated with diverse eukaryotes rather than viruses [9] (**Fig. 7B, 8B**).

### Identification of new putative translation-associated proteins

To explore the function of uncharacterized viral orthologs, we used the detection of distant sequence homology based on tertiary structural information. Interestingly, there were at least four new putative translation-associated proteins (**Table 2**). The predicted structure of MarFTME_090 with high confidence consisting of nine helixes was partially similar to *Saccharomyces* GCN1, which had overlapping binding sites on the ribosome with EF3 [13] (**Fig. 9A, 13**). The partial structure of MarFTME_180 was similar to *Leishmania* zinc-ribbon domain-containing protein, whereas, MarFTME_215 and MarFTME_498 were structurally similar to the whole and part of *Trypanosoma* zinc-ribbon domain-containing protein (DUF4379), respectively (**Fig. 9, 13**). Both zinc-ribbon domain-containing proteins were components of mitochondrial ribosome complexes although their function had not been described [14, 15]. Overall our results indicated that the translation-associated genes of giant viruses might be involved in formation of ribosome complexes.

**Fig. 13.**
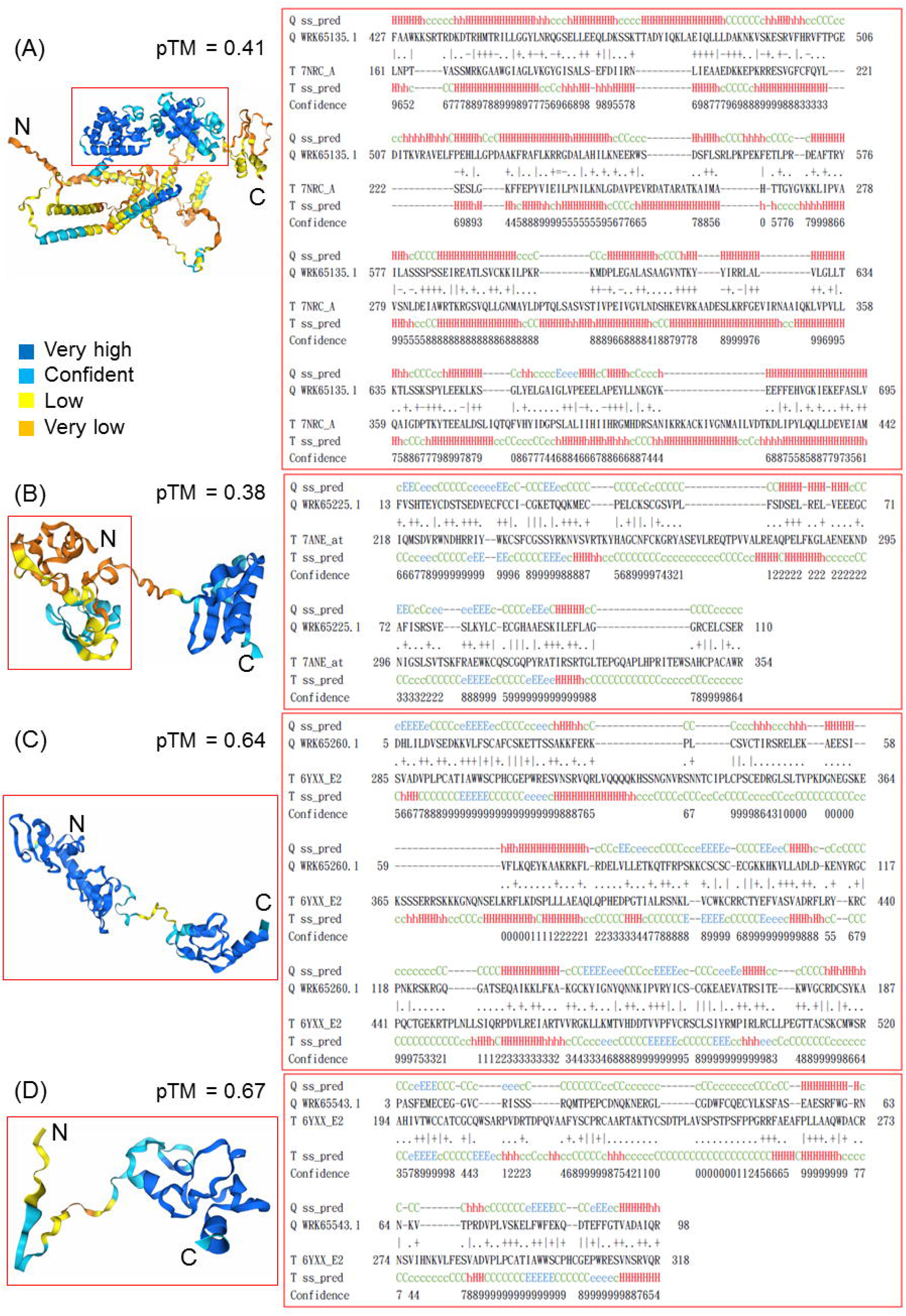
Predicted structures of new viral translation-associated proteins with confidence scores and the multiple sequence alignment between viral protein and templet protein, which were extracted from HHpred. (A) MarFTME_090, (B) MarFTME_180, (C) MarFTME_215 and (D) MarFTME_498. pTM, predicted template modeling score, was derived from a measure called the template modeling score and measured the accuracy of the entire structure. Values higher than 0.8 represented confident high-quality predictions, while values below 0.6 suggested likely a failed prediction.

**TABLE 2.**
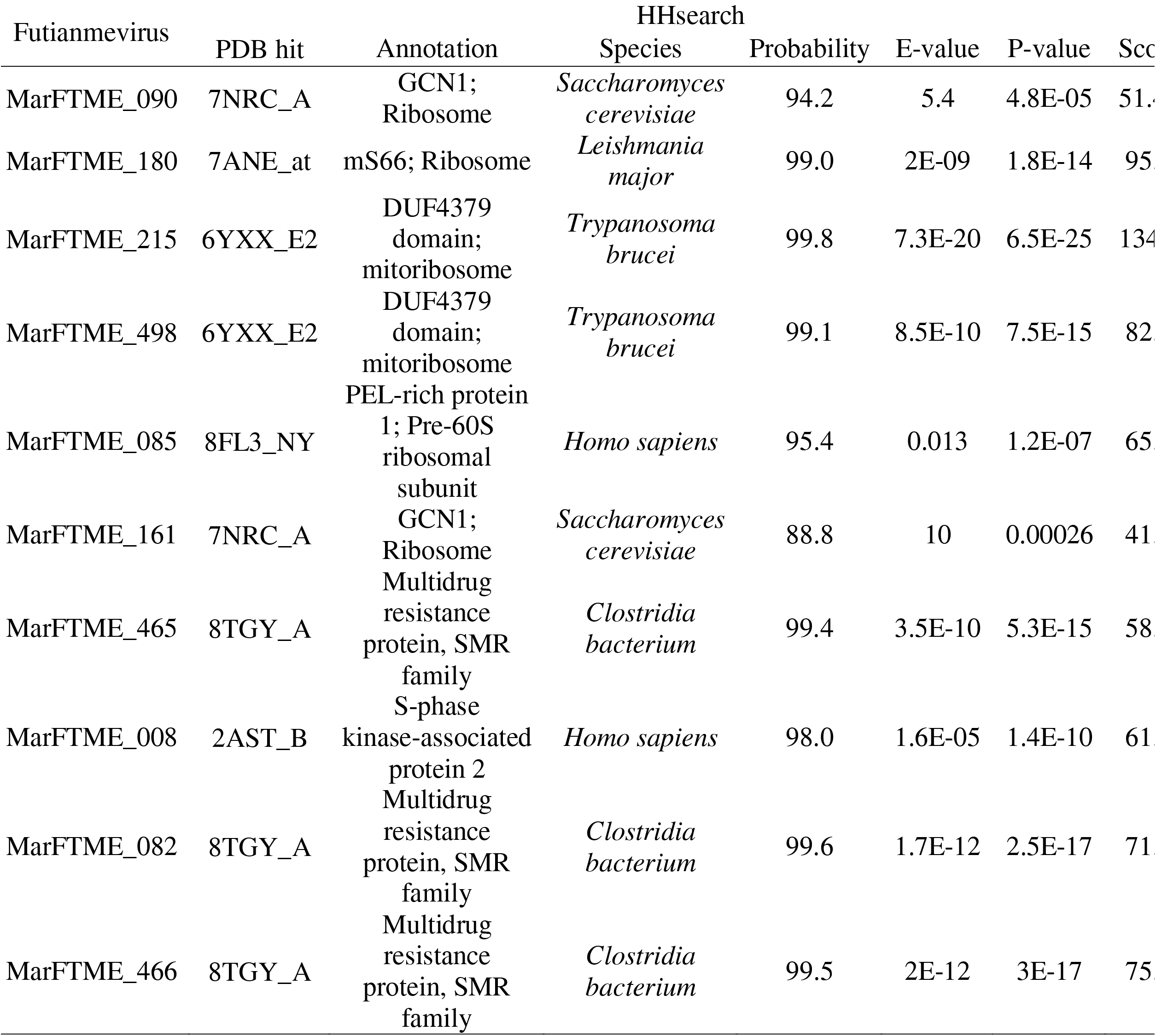
Viral hypothetical proteins annotated based on the best match of tertiary homologue.

## Discussion

As reported in the literature, three treatments, i.e. acid environment (pH 3), high temperature (100 °C), and high concentration of salt (4 M), led to the disruption of mimiviral particle structures, while the alkaline environment or chloroform treatment did not result in the disruption of virions [16]. Although mimivirus and marseillevirus were isolated from the same laboratory acanthamoeba host, here, our results revealed that the biological properties of marseillevirus was different from those of mimivirus.

Marseillevirus was able to form giant vesicles wrapped in membranes derived from host endoplasmic reticulum and the giant vesicles showed better thermal stability than viral particles [7]. Our results revealed the numbers of viral particles released from vesicles varied in the different lineages of marseillevirus and the broad stability range of pH of giant vesicles. The difference of acid/alkaline and thermal tolerance between giant vesicles and viral particles indicated the importance of host membrane for virus fitness under different environments. However, both giant vesicles and viral particles were sensitive to chloroform indicating that vesicles could not protect particles from organic solvents and the lipid component was very important for infection of marseillevirus.

It was hypothesized that the genome repertoire of marseillevirus consisted of genes derived from several distinct sources because amoebae were predators of microbes and “gene melting pots” of microbial evolution [3, 17]. Notably, the viral genes for histone, as well as four components of translation system were hypothesized to be acquired from amoeba or other protists [3]. However, unlike histone proteins, which were identified in all isolated marseilleviruses and proved to be irreplaceable for virus replication [18], translation-associated protein genes were absent in our isolated marseilleviruses. Especially, we isolated for the first time, to our knowledge, a novel marseillevirus lacking EF1A. Considering the phylogenic positions of futianmevirus / futianmfvirus within *Marseilleviridae* and viral EF1A / eRF1 within life domains, our results supported the piecemeal acquisition of translation-associated genes by marseilleviruses [3, 9].

Structural homology facilitated expanding the annotation of previously uncharacterized proteins, sequences of which were orphan in databases and no function assigned. With new protein structures resolved, we identified at least four putative ribosome components in marseilleviruses (MarFTME_090, MarFTME_180, MarFTME_215 and MarFTME_498). The predicted structure of them were similar to the *Saccharomyces* GCN1, *Leishmania* and *Trypanosoma* zinc-ribbon domain-containing protein indicating their roles in translation system, although the function of protozoan zinc-ribbon domain-containing proteins had not been described [14, 15]. The ortholog of MarFTME_090 was found in the proteome of purified virions of Marseillevirs str.T19, namely MAR_ORF226 (YP_003406966.1) suggesting its involvement in early stage of infection [3].

In conclusion, we demonstrated that the four novel diverse marseilleviruses isolated from saltwater samples of mangrove, two of which, futianmevirus and futianmfvirus, were grouped into lineage A, and other two, dashavirus str. E and xiwanvirus, were in lineage B. On one hand, viral particles were halotolerant and acid-tolerant. On the other hand, particles were sensitive to chloroform, highly thermal or extremely acid/ alkaline environment. Additionally, our comparative genomics, phylogenetic analysis, and protein structure comparison revealed diverse translation-associated gene sets in marseilleviruses and novel putative viral components of ribosome complexes.

## Materials and Methods

### Marseillevirus isolation and purification

*Acanthamoeba castellanii* str. Neff (ATCC30010™) were cultivated in peptone-yeast extract with glucose medium (PYG) as described in the literature at 27 °C for 2 days and then the medium was replaced with Page’s amoeba saline (PAS) containing antibiotic cocktail [21].

Saltwater samples were collected from the Futian mangrove reserves, the Xiwan mangrove park and the Dasha river in Shenzhen, Guangdong province, China. The protocols of sample processing, marseillevirus isolation, cloning and titration as well as recipes of solutions had been described in previous works [22].

### Infectivity assay

The infectivity assays were performed by inoculating amoeba cells with the Marseillevirus at an MOI (multiplicity of infection) of 1. At 10 m p.i., the PYG medium was removed and the amoeba cells were washed with PAS three times. At time points between 20 to 1,080 m p.i., the infected cells were collected by cell scrapers into 1.5-ml sterile centrifuge tubes, and then frozen and thawed three times. The cells were collected into new tubes and centrifuged at 356 x g, 5 min at room temperature to remove the cell debris before the supernatants were titrated using the method of end-point dilution [21]. To evaluate the burst size of Marseillevirus, the number of amoeba cells in each well was counted using a counting chamber (Sangon, Shanghai, China), and the total viral particles were obtained as described below.

### Release of viral particles from vesicles

To release viral particles from vesicles, the protocol and recipes of solutions were adopted from the literature [7]. The supernatant obtained from amoeba cells infected with Marseillevirus at an MOI of 1 after 48 h p.i. was submitted to centrifugation at 35,250 x g for 60 min and the pellet containing vesicles was collected. The pellet was resuspended in phosphate-buffered saline (PBS) as a mixture of vesicles and viral particles. To collected particles released from vesicles, the supernatant was mixed with a lysis buffer containing 1% NP-40 and 1% sodium deoxycholate (v/v%) at a ratio of 1:1 for 10 min at room temperature, and centrifuged at 35,250 x g for 1 hour to collect pellets and resuspended in PBS.

### Transmission electron microscopy

The amoeba cells infected by viruses at MOI of 1 were harvest by centrifugation at 356 x g for 5 min, and washed with phosphate-buffered saline (PBS) twice. The cells were fixed with 2.5% glutaraldehyde and stained with 2% osmium tetroxide. The stained cells were dehydrated in a series of ethanol of increasing concentrations and embedded in PON-812 (SPI supplies, PA, USA). Embedded amoeba cells were sectioned at 70 nm thickness using an ultra-microtome (Leica, Wetzlar, German), and fished out onto the 150 meshes. The meshes were stained with 2% uranium acetate avoiding light for 8 min and 2.6% lead citrate avoiding CO_2_. After the meshes were dried at room temperature for 12 h, observations were performed using a HT7700 transmission electron microscope (Hitachi, Japan).

### Chloroform sensitivity

The supernatant containing marseillevirus was treated with chloroform resulting in different final concentrations (10% (v/v%)), and rotated at 60 rpm for 1 hour using rotating mixer shaker (Joan Lab, Zhejiang, China) at room temperature. The chloroform was removed by centrifugation at 500 g for 5 min at room temperature and the aqueous phase was transferred to an new 1.5-ml sterile centrifuge tube and incubated for 12 hour at 4°C to remove any remaining chloroform.

### pH stability assays

The purified marseillevirus pellets were resuspended in PBS, which was adjusted to the desired pH, and rotated at 60 rpm for 24 hour using rotating mixer shaker at room temperature. The supernatant was centrifuged at 35,250 x g for 1 hour to collect pellets and resuspended in PBS (pH 7) and allowed to equilibrate for 12 h. As a control, viral particles were also incubated in PBS (pH 7) for 37 hour at room temperature.

### Thermal stability assays

The marseillevirus supernatants were incubated in a BioRad T100 thermal cycler at 50, 55, 60, and 60 °C for 1 hour and allowed to equilibrate at room temperature for 1 hour. As a control, viral supernatants were also incubated for 2 hour at room temperature.

### High Salty Incubation

The purified marseillevirus pellets were resuspended in PBS, which was adjusted to the desired concentration of NaCl, and rotated at 60 rpm for 24 hour using rotating mixer shaker at room temperature. The supernatant was centrifuged at 35,250 x g for 1 hour to collect pellets and resuspended in PBS (C_NaCl_ = 137 mM) and allowed to equilibrate for 12 h. As a control, viral particles were also incubated in PBS (137 mM) for 37 hour at room temperature.

### DNA preparation

Genomic DNA of the cloned strain of marseillevirus was prepared from the supernatant of infected *A.castellanii* using PureLinkTM Genomic DNA Mini Kit (Invitrogen, Waltham, United States) following manufacturer’s protocol. The protocol of DNA extraction from gram-negative bacterial cells was used. Viral DNA was quantified with a NanodropTM 2000/2000c spectrophotometer (Thermo Fisher Scientific, Carlsbad, CA, USA), as well as by fluorometric quantitation with a Qubit® 3.0 Fluorometer (Thermo Fisher Scientific).

### Viral genome sequencing, assembly and annotation

High-quality genomic DNA (2 μg) was used for DNA Illumina sequencing, which was carried out by the BioMarker Technologies Inc. (Beijing, China). Viral DNA was fragmented into 400-500bp using Covaris M220 ultrasonic instrument. The DNA library was prepared using TruSeqTM DNA Sample Prep Kit (Illumina, San Diego, CA, USA), and sequenced on an Illumina HiSeq Xten with paired-end 150 bp sequencing. Illumina raw reads were processed and the filtered Illumina paired-end reads were assembled de novo using metaWRAP v1.3.2 [23] to obtain the draft assembly.

### Phylogenetic analysis

The nucleotide and amino acid sequences of viruses in the family *Marseilleviridae* including Marseillevirus str. T19 (GenBank accession number: NC_013756.1), Cajuinensisvirus (OR991738.1), Cannes8virus (KF261120.1), Melbournevirus (NC_025412.1), Shanghaivirus (MG827395.1), Tokyovirus (NC_030230.1), Kurlavirus (KY073338.1), Quyangvirus (MT663336.1), Lausannevirus (NC_015326.1), Noumeavirus (NC_033775.1), Portmiou virus (KT428292.1), Insectomimevirus (KF527888.1), Tunisfontaine2virus (KF483846.1), Brazilianmarseillevirus (NC_029692.1), Goldenmarseillevirus (NC_031465.1) were retrieved from GenBank. Megavirus baoshanense (MH046811.2) was used as the outgroup.

Orthologue sequences were extracted from the results of Proteinortho v 6.3.1, which were visualized using the script proteinortho_curves [24], using MUSCLE for multi-sequence alignment. Five NCLDV core genes, including major capsid protein, DNA polymerase delta catalytic subunit, D5-like primase-helicase, virus late transcription factor 3, and A32-like packaging ATPase were chosen for building of the phylogenic trees [12]. The amino acid sequences of archaea and eukaryotic EF1A and eRF1 were extracted from the literature [9].

Orthologue sequences were aligned using MUSCLE v 3.8.1551 [31]. In IQ-TREE multicore v 2.1.2 [25], the best evolutionary models were calculated for each core gene with MFP program to be LG models with GAMMA distributed rates of 5 in all cases. Phylogenetic trees were generated using maximum-likelihood method with LG+G model and 1,000 bootstrap replicates. Phylogenetic trees were visualized using Chiplot v 2.6 [26].

### Comparative genomics analysis

The average nucleotide identity (ANI) was calculated by JSpeciesWS v3.8.5 [27] based on BLAST. The alignments of the nucleotide acid sequences were visualized using BRIG v 0.95 [32]. The gene clusters of viral genes were visualized using Chiplot v 2.6 Sanger sequencing was used to verify the existence of EF1A and eRF1 in marseilleviruses. Amplification reactions were performed using 2x Hieff Canace® PCR Master Mix (Yeasen, China) in a GeneAmp 2400 thermal cycler (Perkin-Elmer-Cetus, Norwalk, CT, USA), and amplification products were gel-purified using HiPure Gel Pure DNA Mini Kit (Magen, China) and T/A-cloned using 5min TA/Blunt-Zero Cloning Kit (Vazyme, China). Multiple recombinants were subjected to Sanger sequencing (GeneWiz Technologies Inc., Suzhou, China), and the sequencing results were compared to identify any potential nucleotide variations.

### Prediction of protein structures

The protein domains were identified for translation-associated proteins using a series software as described in previous works [22]. The amino acid sequences of viral proteins were submitted to HHpred and HHlibs webservers using PDB_mmCIF70_8_Mar and UniRef30_2023_02 [28]. The tertiary protein structures were predicted using Alphafold2 [29], and the results were visualized in PyMol v 2.5.5 [30].

## Acknowledgments

The authors were very grateful to Prof. Jiang Zhong of Fudan University for his useful discussions.

## Author contributions

Y.X. designed research; Y.X. and Y.X. performed research; Y.X. and L.W. analyzed data; Y.X., L.W., Y.W., Z.Z. and R.Z. wrote the paper.

The authors declared that the research was conducted in the absence of any commercial or financial relationships that could be construed as a potential conflict of interest.

